# An integrated, pipeline-based approach for cryo-EM structure determination and atomic model refinement in the cloud

**DOI:** 10.1101/246587

**Authors:** Michael A. Cianfrocco, Indrajit Lahiri, Frank DiMaio, Andres E. Leschziner

## Abstract

Access to streamlined computational resources remains a significant bottleneck for new users of cryo-electron microscopy (cryo-EM). To address this, we have built upon our previous work (Cianfrocco & Leschziner 2015) to develop software tools that will submit cryo-EM analysis routines and atomic model building jobs directly to Amazon Web Services (AWS) from a local computer or laptop. These new software tools (“cryoem-cloud-tools”) have incorporated optimal data movement, security, and cost-saving strategies, giving novice users access to complex cryo-EM data processing pipelines. Integrating these tools into the RELION processing pipeline and graphical user interface we determined a 2.2 Å structure of ß-galactosidase in ~55 hours on AWS. We implemented a similar strategy to submit Rosetta atomic model building and refinement to AWS. These software tools dramatically reduce the barrier for entry of new users to cloud computing for cryo-EM and are freely available at <u>*cryoem-tools.cloud*</u>.

## INTRODUCTION

Cryo-electron microscopy (cryo-EM) is a structural biology technique that has undergone rapid growth over the past few years (Nogales 2016). Technical developments in direct electron detection and electron optics in conjunction with improvements in image analysis (Scheres 2014; Punjani et al. 2017) have led to the widespread adoption of cryo-EM as a structural biology technique. Furthermore, the advent of GPU-accelerated cryo-EM structure determination (Punjani et al. 2017; Kimanius et al. 2016) has helped to reduce the overall cost for computing hardware for a single user. While these improvements have helped to spread cryo-EM, it becomes difficult to scale the required hardware to accommodate large cryo-EM facilities that have a large number of users. These facilities have to balance cost with availability of resources: idle computing infrastructure is wasted capital whereas queuing times for compute resources waste personnel salaries. The challenge is how to create a computing facility that is cost-effective while also delivering compute resources on-demand without wait times.

We have previously shown that Amazon Web Services (AWS), the world’s largest cloud computing provider, is a cost-effective resource for cryo-EM structure determination (Cianfrocco & Leschziner 2015). Since this original publication, AWS released GPU-accelerated virtual machines (‘VMs’) (named ‘p2’ and ‘g3’) with 1, 8, or 16 NVIDIA K80 GPUs on p2 or 1, 2, or 4 NVIDIA M60 GPUs on g3, while also reducing prices for data storage on the block storage service (‘S3’) and archival storage (‘Glacier’).

Despite its power, our original work required users to manually deploy AWS resources. To streamline the process, we have developed software tools that allow for the remote management of AWS resources from the local computer of a user. These tools were then combined with the standard suite of cryo-EM software tools MOTIONCOR (Li et al. 2013), MOTIONCOR2 (Zheng et al. 2017), UNBLUR (Grant & Grigorieff 2015), GCTF (Grant & Grigorieff 2015; Zhang 2016), CTFFIND4 (Rohou & Grigorieff 2015), RELION (Scheres 2012) and Rosetta (Wang et al. 2016; Wang et al. 2015), allowing users to submit jobs directly to AWS from their local project directory while syncing results back in real time. In contrast to our previous implementation, we are now using ‘on-demand’ VMs from AWS, which eliminates the risk of users being ‘kicked-off’ due to price changes. Finally, by combining the full RELION pipeline (and associated software) with atomic model building and refinement with Rosetta (Wang et al. 2016; Wang et al. 2015) with AWS, cryoem-cloud-tools provides users with all aspects of cryo-EM structure determination in a single pipeline - from micrograph motion correction to atomic model refinement.

## APPROACH

We realized that the workflow from our original publication (Cianfrocco & Leschziner 2015) was cumbersome, requiring users to interact with AWS resources using complex commands. To streamline this process, we wrote software tools that leverage the capabilities of command-line tools provided by AWS. Then, we incorporated these commands directly into the RELION GUI to allow users to submit RELION jobs directly to AWS (Figure 1).

**Figure 1.**
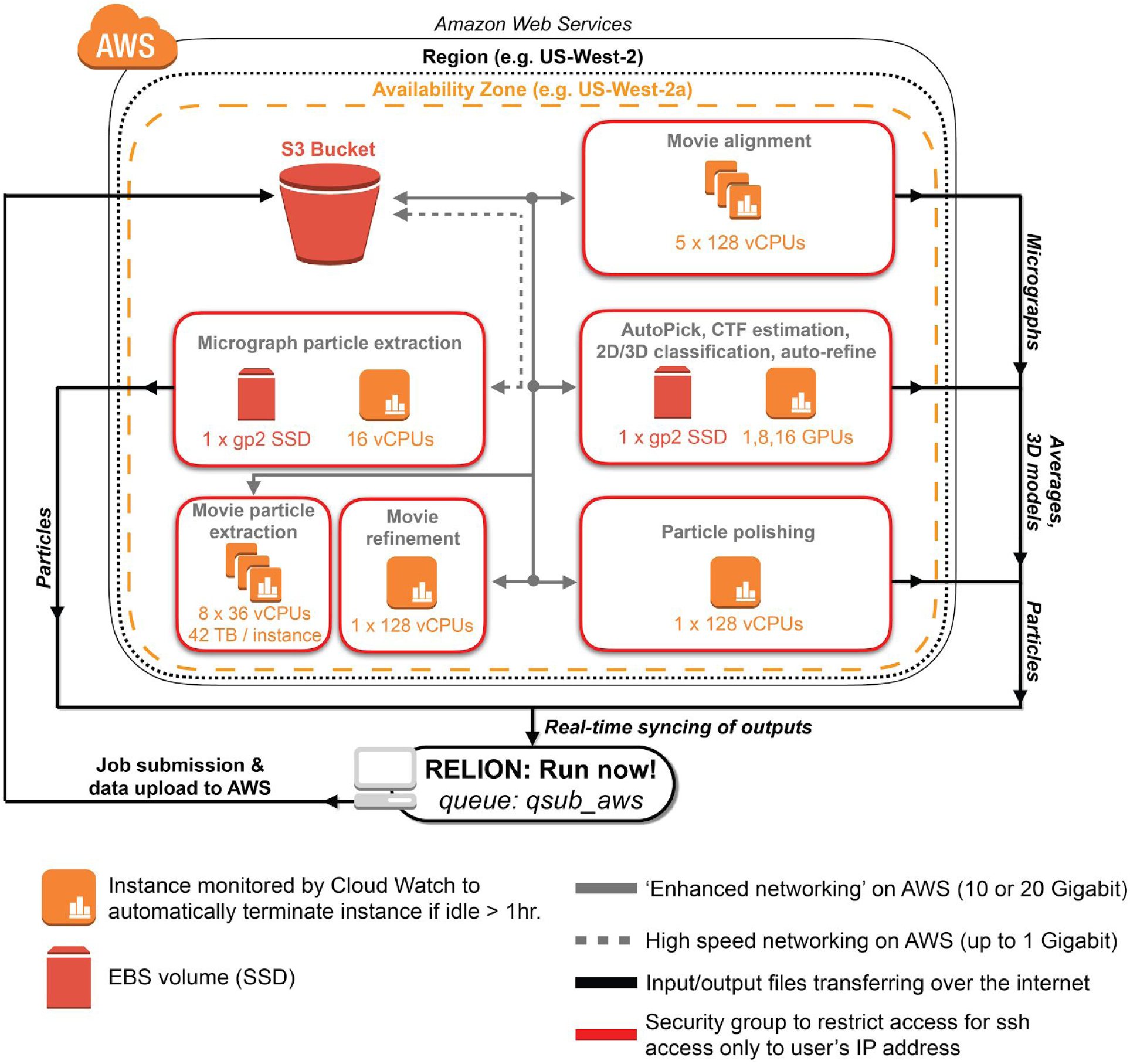
AWS architecture for cryo-EM data processing with RELION. Shown is aschematic of AWS resources deployed by cryoem-cloud-tools through the program ‘qsub_aws’. For all job types shown, the software places VMs within security groups that restrict access to the IP address of the end-user. Within a security group, the software determines the appropriate VM and storage choices, using S3 as a distribution point between local and AWS resources.

The overall approach takes advantage of the cluster submission feature of RELION by providing users with a new submission command (‘qsub_aws’) to do the following: 1) identify the type of RELION job, 2) upload data to AWS block storage (S3), 3) start VM(s) required for the task, 4) download data from S3 to VM, 5) Run RELION commands, 6) Sync output results back to the local machine in real time, and 7) Turn off machines when finished (or if an error is detected). As shown in Figure 1, we implemented job type-dependent data processing strategies for RELION analysis routines. This means that GPU-accelerated steps (Movie alignment, Auto Pick, CTF estimation, 2D/3D classification, auto-refine) are run on VMs with GPUs (p2 VMs), whereas CPU-based steps are run on VMs with 16 or 128 virtual CPUs (vCPUs) (See (Cianfrocco & Leschziner 2015) for a detailed discussion of vCPUs vs. CPUs).

In building this software, we are providing users with workflows that have been optimized for data transfer and computing. For instance, all data is first uploaded into AWS’s S3 ‘buckets’. This allows for fast uploads (up to 300 MB/sec) and also for cost-effective storage of data in-between analysis routines. Storing data on S3 between RELION runs removes the latency that results from re-uploading the same data multiple times. Next, we implemented data storage policies that allow for high input/output tasks and large dataset sizes, which included 42 terabyte drives for movie particle extraction on d2 VMs. Finally, for computational tasks that can be distributed (Movie alignment and Movie particle extraction), we boot up and manage multiple VMs in parallel to finish analysis routines quickly.

## RESULTS & DISCUSSION

To assess the performance of our approach, we compared processing times for the determination of a 2.2 Å ß-galactosidase structure (Bartesaghi et al. 2015) that was recently solved using a stand-alone GPU workstation (Kimanius et al. 2016). While the comparison is testing very different computing environments, we chose it because many new cryo-EM users are purchasing stand-alone GPU workstations and we wanted to compare performance relative to AWS.

Using our integrated AWS software tools in the RELION GUI and launching all RELION analysis commands remotely, we were able to determine a 2.2 Å structure in 54.5 hours on AWS (Figure 2A & 2B, Figure 2 - Supplement 1), which is 2X faster than a standalone GPU workstation (Figure 2C). These processing times also included the time required for movement of data into and between resources on AWS, thus reflecting the full processing times experienced by a user. For GPU-accelerated RELION processing steps, VMs with 8 GPUs (p2.8xlarge) performed equally well or slightly faster than a 4 GPU workstation (Figure 2D). This likely results from faster GPUs in the workstation (NVIDIA GTX1070: 1683 MHz clock speed) compared to those on AWS (NVIDIA K80: 875 MHz clock speed). Expectedly, the largest improvements in time saved were seen in steps that could be distributed across multiple VMs (Movie alignment and Movie particle extraction) (Figure 2D). For these processes, we were able to select VMs that were appropriate for the process - GPU machines for movie alignment (p2.8×large), large storage arrays for movie particle extraction (d2.8×large), and high vCPU numbers for movie refinement and polishing (×1.32×large: 128 vCPUs).

**Figure 2.**
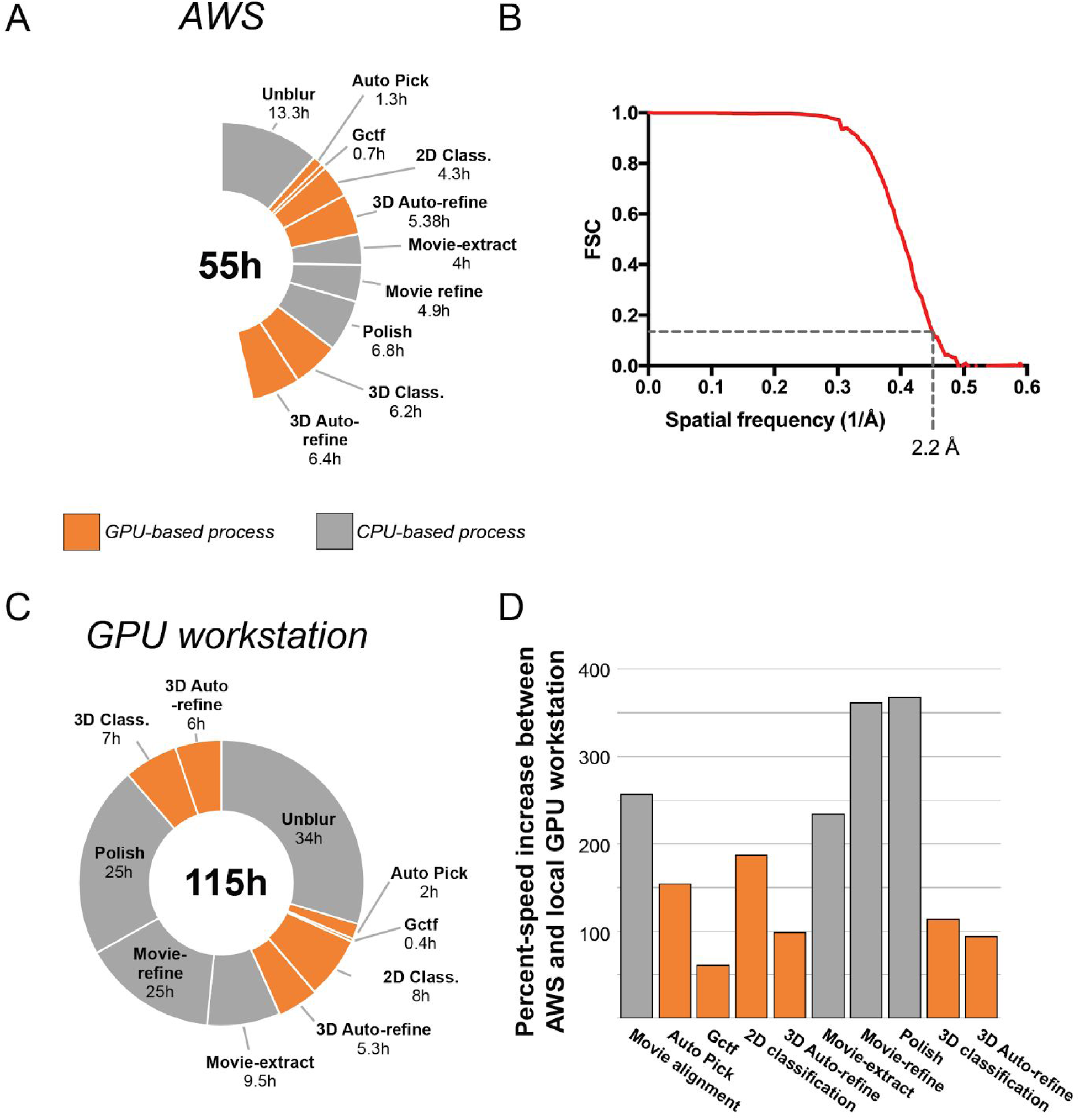
Performance of AWS vs. local GPU workstation. Processing times (A) and FSCcurve (B) for the determination of a 2.2 Å ß-galactosidase structure on AWS. (C) Processing times from the determination of 2.2 Å ß-galactosidase structure on GPU workstation (Kimanius et al. 2016). (D) Comparison of percent speed-up increases between AWS and a GPU workstation.

In order to build the atomic model for ß-galactosidase into this density, we used the molecular modelling program Rosetta (Wang et al. 2016; Wang et al. 2015). As modelling software, Rosetta needs CPU computing clusters because its sampling of hundreds of atomic models relative to the cryo-EM density requires a dedicated CPU for each model. Therefore, we incorporated Rosetta tools for model building and refinement into our AWS-based pipeline, allowing users to submit a Rosetta refinement to AWS from their local computer or laptop (Figure 3A). By distributing the Rosetta refinement over multiple VMs on AWS, each with 36 vCPUs (c4.8×large), we were able to generate 200 models using RosettaCM and Rosetta *FastRelax* 6.1 hours on AWS, a speedup of about 7X over a single workstation with 16 processors (41.8 hours) (Figure 3B). The resulting model showed good agreement with the density, where the r.m.s.d for the top 10 Rosetta models was < 0.5 Å. (Figure 3C & Figure 3D, Figure 3 - Supplement 1).

**Figure 3.**
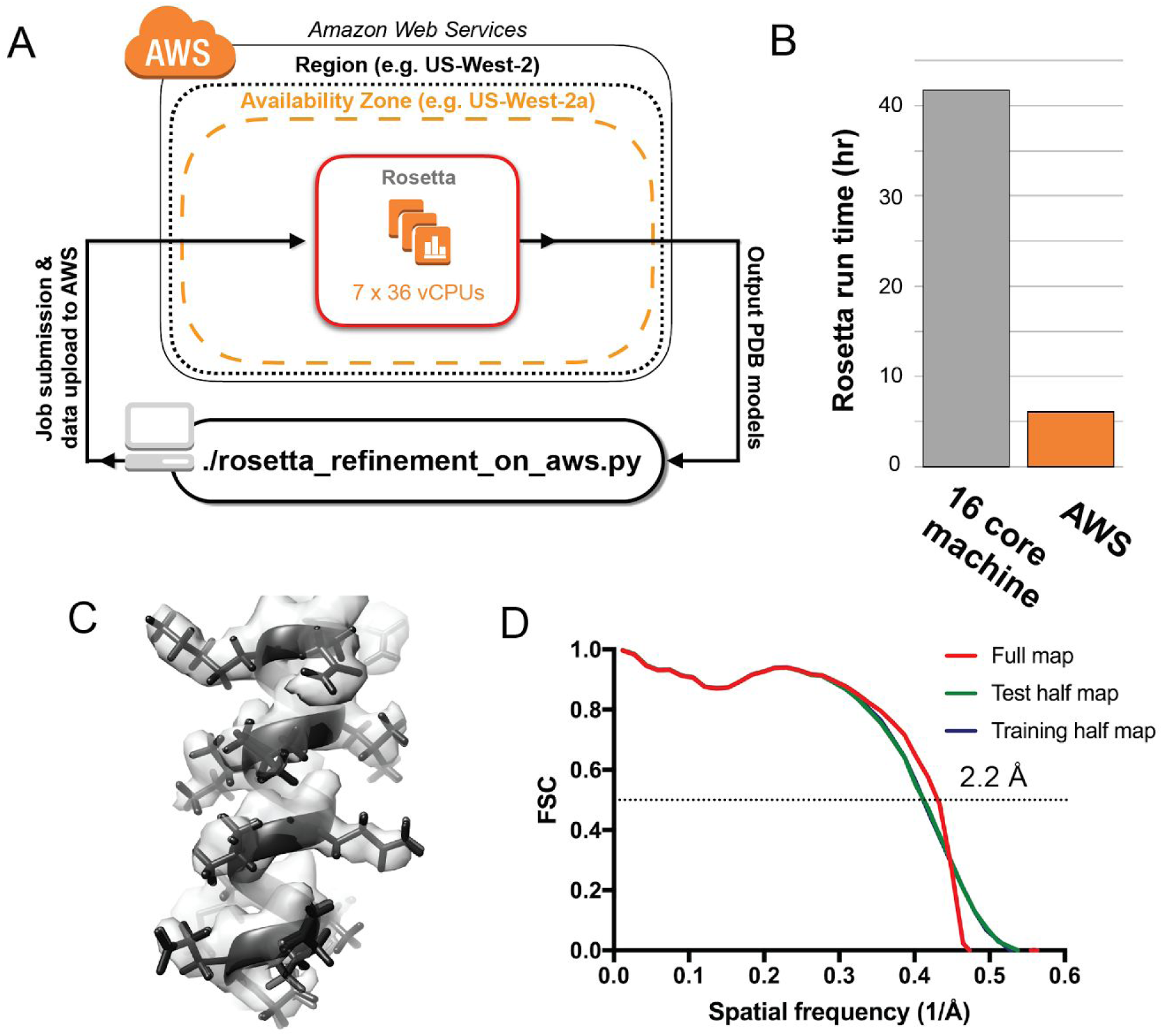
Rosetta atomic model refinement in the cloud. AWS architecture for running Rosetta model refinement across multiple VMs. (B) Run time comparisons between a local workstation (16 cores) and AWS (252 vCPUs). (C) Representative region of the cryo-EM map with the top five atomic models built by Rosetta *FastRelax* (D) FSC curves between the best atomic model from *FastRelax* and the cryo-EM map of ß-galactosidase.

The cost for determining a 2.2 Å structure using RELION and building an atomic model with Rosetta, both using AWS, was $1,426 USD (Figure 2 - Supplement 1). This cost represents both storage and computing on AWS, with the top three expenditures (71% of the total) coming from 30 terabytes of data storage on AWS S3 ($690.00), Movie particle extraction ($179.73), and Movie alignment ($146.72) (Figure 2 - Supplement 1).

A typical user may not use AWS for those computation-intensive steps (movie processing) that accounted for most of the cost of solving this structure. If we consider a scenario where a user performs movie alignment locally and does not perform Movie refinement or Polishing, but submits all other jobs to AWS (AutoPick, CTF estimation, 2D & 3D classification, 3D auto-refine, and Rosetta), the total cost would be $260.41 USD. Since this latter scenario appears to be more prevalent with the advent of per-particle tracking and dose weighting during pre-processing of data (Zheng et al. 2017; Rubinstein & Brubaker 2015), AWS offers users with an accessible computational resource for cryo-EM.

This approach for cryo-EM data analysis has the potential to benefit many different types of cryo-EM users. Since this software package integrates directly into a user interface, individual users will have the option to perform multiple analysis routines from a single workstation by pushing additional jobs to AWS instead of waiting to run them sequentially on a local GPU workstation. For research teams, this software provides ‘burstable’ processing power, ensuring that data processing does not become rate-limiting ahead of grant and manuscript deadlines. Finally, this software can have a significant impact on cryo-EM facilities with a large user base. Given the scale of AWS, a cryo-EM facility could not only provide many users with access to microscopes but also allow those users to push cryo-EM jobs to AWS without having to accommodate their computing needs locally.

## Data Accessibility

Information and tutorials related to using the software presented here are available at <u>*cryoem-tools.cloud*</u>. All software is freely available at Github: https://github.com/cianfrocco-lab/cryoem-cloud-tools. The ß-galactosidase cryo-EM structure can be accessed at EMD XXXX and PDB XXXX.

## ACKNOWLEDGEMENTS

We would like to thank all of the members of the Leschziner lab at UC San Diego for helping to test and debug commands to run on AWS. M.A.C. was an HHMI Fellow of the Damon Runyon Cancer Research Foundation. A.E.L. is supported by grant R01GM107214 from the National Institutes of Health and F.D. is supported by grant R01GM123089 from the National Institutes of Health.

## METHODS

### Integrating cryoem-cloud-tools into the RELION GUI

The overall strategy for users accessing cryoem-cloud-tools from the RELION GUI utilized the cluster submission of RELION. When users submit jobs to a cluster, they indicate the submission command directly into the RELION GUI (e.g. ‘qsub’). Within this framework, we built cryoem-cloud-tools to be specified directly from the GUI using a python program named ‘qsub_aws.’ This program will automatically determine the type of RELION command that needs to be run and determine the AWS resources required to execute the task. This approach does not require users to compile RELION using cryoem-cloud-tools; instead cryoem-cloud-tools is a software extension for RELION to submit jobs to AWS.

### ß-galactosidase image processing

To replicate the published work on ß-galactosidase (Kimanius et al. 2016; Bartesaghi et al. 2015), we used an almost identical processing strategy. A summary of processing times, VM types, and costs can be found in Figure 2 - Supplement 1. All VMs were ‘on-demand’, which means that we paid full price and did not risk being ‘kicked off’ by being outbid due to spot price markets. We uploaded 1536 7676 × 7420 pixels super-resolution movies of ß-galactosidase (EMPIAR 10061) (Bartesaghi et al. 2015) to AWS and aligned them using Unblur (Grant & Grigorieff 2015) on 5 × ×1.32×large instances. From our data servers at UCSD, we were able to achieve ~350 MB/sec upload speeds to S3 using multi-file uploads with ‘rclone’. Gctf (Zhang 2016) was used to estimate the CTF of the aligned micrographs on a single p2.8xlarge VM (8 GPUs). Then, 138,901 particles were picked using GPU-accelerated AutoPick on a single p2.8xlarge VM and extracted at a pixel size of 1.274 Å (binned by 4 from the original data) in a box size of 192 × 192 pixels on a single m4.4×large VM (16 vCPUs). This stack of particles was subjected to 2D classification into 200 classes over 25 iterations on a p2.16xlarge VM (16 GPUs). election of the best class averages resulted in a stack of 119,443 particles that were then re-extracted at a pixel size of 0.637 Å in a box size of 384 × 384 pixels on a m4.4×large VM. These particles were refined with PDB 3I3E (Dugdale et al. 2010) as the initial model using auto-refine to a resolution of 3.5 Å (unmasked) on a single p2.8×large VM. These refined coordinates were used for Movie particle extraction on 8 × d2.8×large VMs (36 vCPUs and 48 Terabytes on each VM) and Movie refinement on a single x1.32xlarge VM (128 vCPUs) with a running average of 7 movie frames and a standard deviation of 2 pixels on particle translations. These particles were subjected to Polishing on a single ×1.32×large VM, yielding an unmasked resolution of 3.3 Å, after which they were used for 3D classification into 8 classes over 25 iterations using an angular step of 7.5 degrees on a single p2.8×large VM. From the 4 best classes, 106,237 particles were used for 3D auto-refine on a single p2.8×large instance to obtain a final, post-processed structure at 2.2 Å, as previously reported (Kimanius et al. 2016; Bartesaghi et al. 2015).

### Atomic model building with Rosetta on AWS

We extended cryoem-cloud-tools to allow users to build atomic models into cryo-EM maps using Rosetta, specifically RosettaCM and Rosetta’s *FastRelax* protocols. We ran these protocols on c4.8xlarge instances with a single solution requested per vCPU. Using this method we generated atomic models for the 2.2 Å ß-galactosidase map determined on AWS. We used atomic coordinates of 1JZ7 chain A as the starting model for the asymmetric unit of the ß-galactosidase map and generated the initial aligned reference structure using *rosetta_refinement_on_aws.py* routine from cryoem-cloud-tools. Following this step, we generated the symmetry definition file for Rosetta describing the D2 symmetry of ß-galactosidase in the context of 1JZ7 using the script *rosetta_prepare_symmfile.py*. All these initial steps were carried out on t2.micro instances. We used the initial reference structure and the symmetry definition file as input and used RosettaCM to generate 200 output models. RosettaCM was run using using *rosetta_refinement_on_aws.py* routine running on 10 x c4.8xlarge instances with 20 models per instance. The best model in terms of Rosetta energy (including fit-to-density energy) was used as an input for a final refinement with Rosetta’s *FastRelax*. We generated 8 models from *FastRelax* using one of the two half maps generated during refinement (training half map) lowpass filtered to a resolution of 2.24 Å and sharpened with a B-factor of -49.52. To estimate overfitting, FSC_work_(FSC curve between the refined model work and the training half map) and and FSC_work_(FSC curve between the refined model and the other free half map generated during refinement, the test half map) were compared and the the spatial frequency at which the FSC value was 0.5 was 1/2.4 Å^-1^ in both cases. The FSC curves were calculated in Rosetta and the plots were made using GraphPad Prism (GraphPad software). The best model in terms of Rosetta energy and model geometry (as determined by MolProbity) was selected as the final atomic model for the ß-galactosidase map.

## Figures

**Figure 2 - Supplement 1 -.**
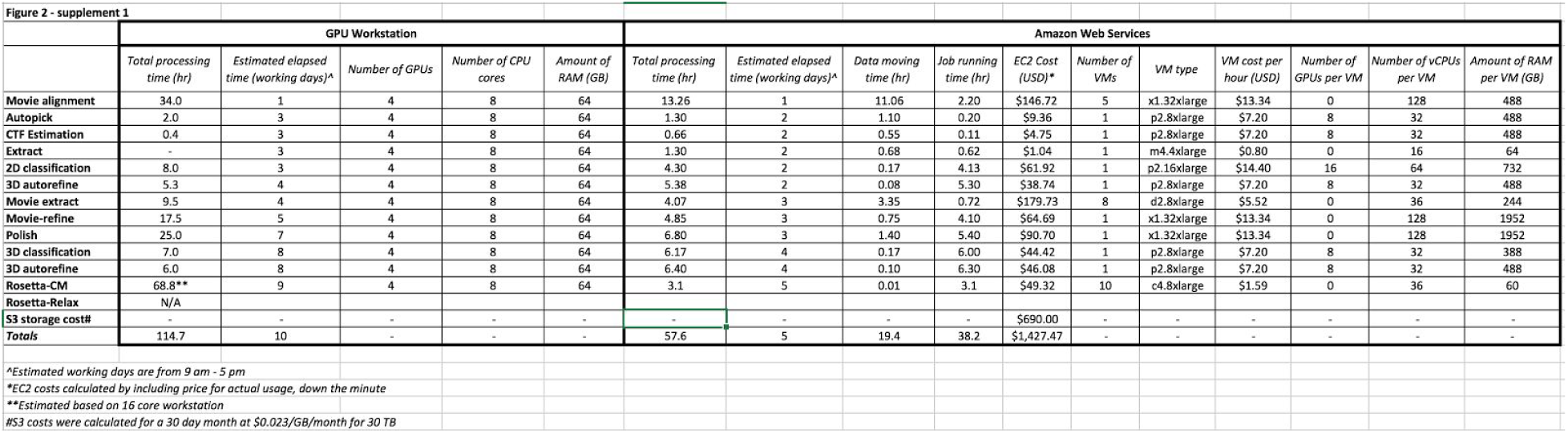
Summary of processing times and costs associated with ß-galactosidase structure determination on AWS compared with a local GPU workstation.

**Figure 3 - Supplement 1 -.**
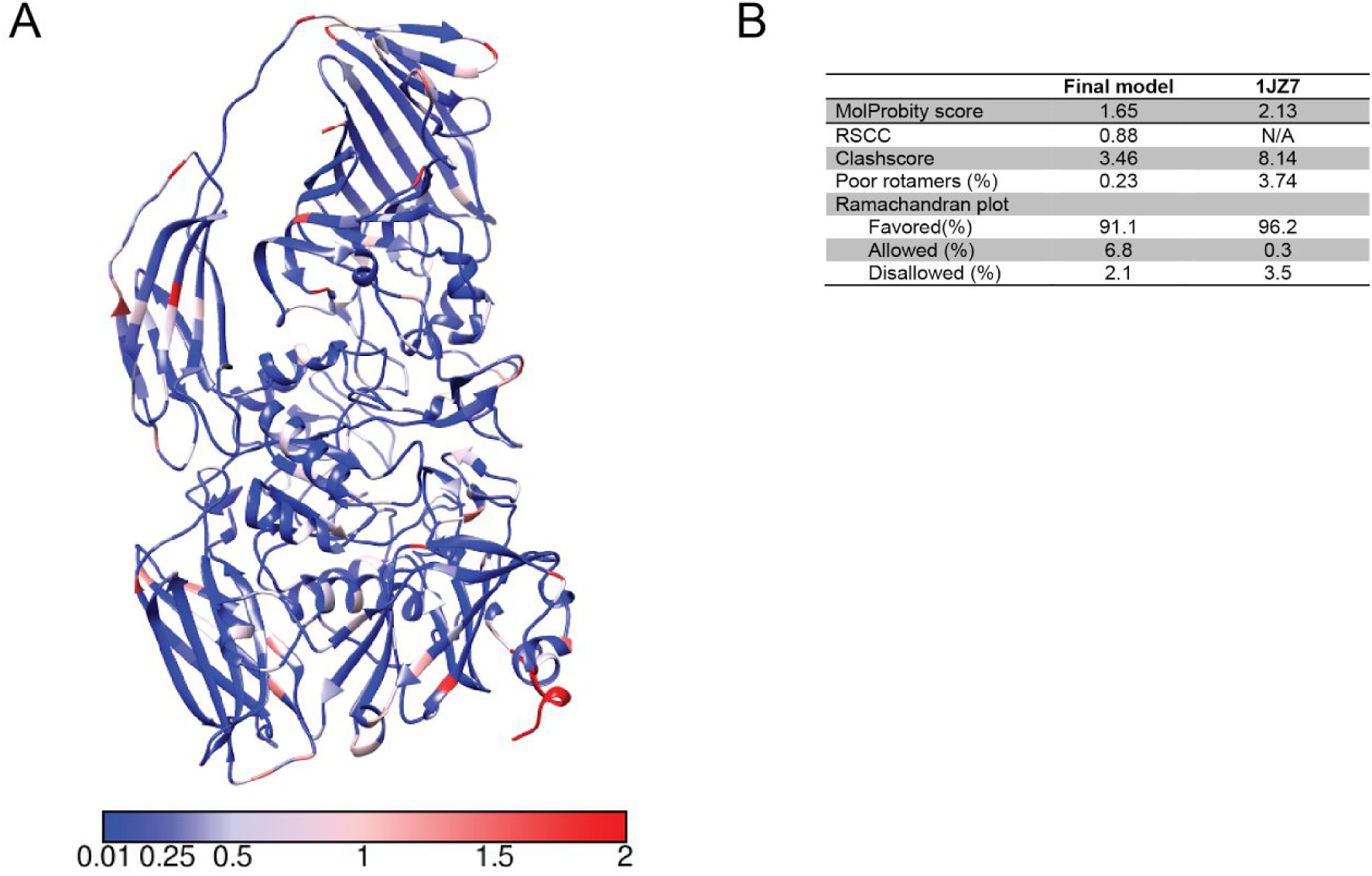
Rosetta modeling statistics. The RosettaCM model used as input for Rosetta *FastRelax* colored by the the all atom r.m.s.d. value of the top ten RosettaCM models (based on Rosetta energy). Units of scale are Å One of the four asymmetric units is shown. (B) Table summarizing the model validation statistics determined by MolProbity for the final atomic model. For comparison the model statistics of the starting model, 1JZ7 is shown.

